# High-speed MRI recordings of eyeball lifting, retraction and compression during blinks

**DOI:** 10.1101/2022.05.11.491482

**Authors:** Johannes Kirchner, Tamara Watson, Jochen Bauer, Markus Lappe

## Abstract

Blinks occur frequently in normal life and have increasingly been linked to perceptual and cognitive effects. However, the oculomechanics of blink-related eye movements are much less researched than other types of eye movements. While it has been observed that the eye is being pulled back into its socket during a blink, possibly due to co-contraction of extraocular muscles, this elusive eye motion has not been studied in detail due to the technical difficulties that go along with a closed eyelid. Here we use dynamic magnetic resonance imaging (MRI) to obtain videos of this motion and analyse the kinematics with the recently developed MREyeTrack algorithm. We show that the eye is not only retracted but also lifted up during a blink. For some participants we observed eyeball lifting by up to 3 mm, far exceeding the amount of translation believed to occur during natural eye movements. Slow blinks can be accompanied by large tonic rotations of up to 15°. Furthermore, we collected evidence that the co-contraction of extraocular muscles leads to a slight compression of the eyeball. These findings demonstrate the surprising complexity of ocular motility and offer new opportunities to study orbital mechanics in health and disease.

## Introduction

Blinks are a fundamental aspect of the human visual system, occurring around 4 times per minute [1]. Their primary function is maintaining a healthy cornea by wetting and removal of irritants from its surface [2], but beyond that blinks are involved in a variety of tasks. The brief disruption of vision caused by blinks goes mostly unnoticed due to active visual suppression [3, 4] and is used to readjust the oculomotor system [5–7]. Blinks also play a crucial role in allocating attention and have, for example, been found to occur strategically timed in response to environmental tasks [8] and at key break points when watching videos [9]. Since damage to the eye from mechanical interference might easily lead to blindness, many species in the animal kingdom have a mechanism protecting the eye from harm. Giant guitarfish, like many fish and amphibians, cannot cover the eye with their lids, so they let the entire eye sink completely into their head by contracting their eye muscles [10]. Mammals like guinea pigs and rabbits can cover their eyes by blinking, but this is also accompanied by retracting their eyes [11]. In order to do so, they have a specific muscle assigned to this, the retractor bulbi.

Humans do not have a retractor bulbi muscle, but their eyes nevertheless retract while blinking [2, 11–13]. This retraction is believed to be performed by co-contraction of several extraocular muscles [11, 14, 15]. The exact innervation pattern of this co-contraction has so far not been measured in humans. In rabbits, EMG recordings have shown that all extraocular muscles except the superior oblique contract, but it has been argued that the contraction of inferior and superior rectus would be sufficient to explain both retraction and rotational motion during blinks in humans [15]. The exact extent of the blink-related eye movement in humans has a somewhat controversial history. It was first described by Charles Bell in the early 19th century as a large upwards and outwards rotation. Reports of a more diverse motion followed but an upwards rotation was still found in the large majority of participants during visual observation when forcefully holding the lid open [16]. This upwards rotation found its way into neurology and ophthalmology textbooks as Bell’s phenomenon and was assumed to take place during normal blinks as well. Modern studies using scleral search coils could not confirm this finding. Instead they showed that the eye rotated slightly downwards by 2-5° [14, 15]. Around the same time evidence for eyeball retraction appeared [2, 11], which suggested that the blink-related eye movement is mainly a translational motion which is accompanied by some amount of incidental rotation [14, 15]. The discrepancy with reports from visual observation could be somewhat resolved by the observation of large, tonic rotations for longer, forceful blinks but these could be either up- or downwards [14]. Some of the controversy remains and in particular detailed accounts of the translational motion during blinks are still missing.

Little is known about eyeball translation in general. Conventional eye tracking techniques like search coils, EOG or video-based methods do not measure translational motion and operate under the assumption that eye movements are purely rotational, yet small translations have been observed as a function of static gaze direction [17, 18], of changes in fixation distance [19] and during saccades [20]. Small translations probably do not have large perceptual consequences for the visual system, but they could still be of importance for the ocular mechanics due to changes in torque lever arms of the extraocular muscles [17]. Abnormal eyeball translation has also been observed in strabismus [21], with retraction even being the main syndrome in some rare cases [22, 23]. Blinks are unique among natural eye movements in that their main motion component is translation and not rotation.

In this study, we used high-speed dynamic MRI to record the blink-related eye movement from eleven participants at a temporal resolution of 52-54 ms. Dynamic MRI sequences have been successfully used to track the motion of human organs like the liver [24], pancreas [25] or eye [13]. MRI allows to image an entire cross-section of the eye, which has the advantage of measuring eye movements even when the lid is closed and visualising displacements and deformations of the whole eyeball. Eyeball kinematics, in particular the translational motion, were estimated from single-slice data using the recently developed MREyeTrack algorithm [13]. Participants were instructed to fixate a dot which was presented at central gaze position and to make voluntary blinks of various durations. This allowed us to analyse the blink-related eye movement both during natural, short blinks as well as during prolonged periods of eye closure. We also investigated whether co-contraction might be accompanied by eyeball compression by tracking changes in eyeball diameter during blinks.

## Methods

### Participants

Eleven healthy participants (P1-P11, age 23-49, one female, ten males) took part in this study. Participants gave informed consent and all procedures were approved by the ethics committee of the Department of Psychology and Sports Science of the University of Münster.

### Experimental Setup

Participants were examined by a 3T Philips Achieva MRI scanner (Philips Medical Systems, Best, The Netherlands). Instructions and visual stimuli were displayed using a back-projection monitor which was placed at a total viewing distance of 108 cm. First, we acquired 3D T2 weighted MR data of the entire head (matrix = 256 x 256 x 250, FOV = 250 x 250 x 250 mm, voxel size = 0.98 x 0.98 x 1.00 mm, TE = 225 ms, TR = 2500 ms, slice thickness = 1 mm, flip angle = 90°, scan duration = 232.5 s), which was used as a reference for planning the following dynamic single-slice scans. Eyeball motion was captured using a balanced steady-state free precession (bSSFP) sequence either in the axial plane (matrix = 224 x 224, FOV = 200 x 200, voxel size = 0.89 x 0.89 mm, TE = 1.28 ms, TR = 2.56 ms, slice thickness = 3 mm, flip angle = 45°, 1020 dynamic scans, total scan duration = 56.7s) at a temporal resolution of 55.6 ms or in the sagittal plane (matrix = 224 x 224, FOV = 200 x 200, voxel size = 0.89 x 0.89 mm, TE = 1.24 ms, TR = 2.47 ms, slice thickness = 3 mm, flip angle = 45°, 1020 dynamic scans, total scan duration = 54.2s) at a temporal resolution of 53.1 ms. In both dynamic scan series a k-t BLAST factor of 5 were used to accelerate the acquisition per slice.

### Experimental Protocol

During the initial T2 weighed 3D data acquisition, participants were instructed to fixate a black dot of 0.8° diameter on a grey background, so that eyeball motion could be minimized. During the dynamic bSSFP scans, participants had to execute one of the three tasks *Short Blink, Slow Blink* or *Eye Closure*, each while fixating the same black dot at center position. For the *Short Blink* task, participants were instructed to aim for a short, natural blink. Next, we asked participant to make their blinks a bit longer than usual during the *Slow Blink* task. Finally, participants were instructed to repeatedly close their eyes for a full second during the *Eye Closure* task. Each participant had to perform each task twice, once while recording the eye motion in the axial plane and once in the sagittal plane. We continuously monitored the imaging slice position between different acquisitions and adjusted the slice position if necessary to ensure that the lens was visible. For stimuli presentation and data analysis we used MATLAB (The MathWorks, Natick, MA, USA) with the Psychophysics Toolbox [26].

### Data Analysis

#### Preprocessing

Image intensities for all MR data were rescaled such that the intensities around eyeball center corresponded to a value of one. This was done in order to make the metrics of the subsequently used segmentation algorithm comparable across participants and sequences. Furthermore, head motion in the dynamic bSSFP scans was estimated using an efficient sub-pixel image registration by cross-correlation algorithm [27].

#### Eye Tracking

Eye motion in the dynamic bSSFP scans were quantified using the MREyeTrack algorithm described in Kirchner et al. [13], which allows to measure both rotational and translational motion components. The algorithm produces a segmentation of sclera, lens and cornea by finding the optimal projection of a 3D eyeball model for each image. We introduced the following minor modifications to the algorithm. Some of the MR recordings had artefacts in the anterior segment of the eyeball, in particular around the cornea. Therefore, we introduced weights to the individual components of the energy functional to be minimised in MREyeTrack. The original functional *E* = *E*_sclera_ + *E*_lens_ + *E*_cornea_ consisted of equal contributions from sclera, lens & cornea, which we changed to *E* = *E*_sclera_ + 0.5 * *E*_lens_ + 0.25 * *E*_cornea_ to make it more robust to artefacts in the anterior segment of the eye. Also, we constricted out-of-plane translation to a maximum of 3 mm and out-of-plane rotation completely. The obtained eye motion time series data was upsampled to a 2 ms time interval via linear interpolation and then smoothed using a Savitzky-Golay filter of 2nd order with a window of 100 ms [28] for further analysis.

#### Blink detection

Simultaneous measurements of pupil area and eyeball translation have shown that blinks are always accompanied by translation [13]. Therefore, we defined and detected blinks in our data based on the measure of eyeball translation along the anterior-posterior axis, i.e. retraction, which is present in both sagittal and axial MR recordings. The eye retracts at relatively high velocity during lid closure, so blinks were coarsely located by finding samples where the retraction velocity exceeded 2 mm/s. Blink onset was then defined as the first sample the velocity exceeded 1 mm/s. Accordingly, blink offset was determined when the eye propelled back and subceeded a velocity of 1 mm/s. We required blinks to have a minimum duration of 100 ms and a minimum retraction of 0.3 mm.

#### Eyeball compression

The MREyeTrack algorithm works with a rigid eyeball model and therefore does not incorporate eyeball compression, which could hypothetically occur due to the co-contraction of several extraocular muscles during a blink. To investigate this, we calculated the longitudinal and transversal diameter of the eyeball during short blinks. The longitudinal diameter was defined along the axis running through both eyeball and lens center, while the transversal eyeball diameter was defined to be perpendicular to that. Borders of these diameters were defined as the locations where the image intensities fell below a value of 0.5. Only the posterior border required a different threshold, because the tissue behind it was often of higher intensity. Therefore, the posterior border was determined as the average intensity between 3 mm before and after the MREyeTrack segmentation.

#### Data exclusion criteria

MRI is prone to susceptibility artefacts at natural interfaces like that of air and tissue, which can manifest themselves as a local distortion of the magnetic field and subsequent loss of anatomic structure in the image. MR recordings of some participants showed a loss of anatomic structure at the anterior segment of the eye. If these artefacts are small, the MREyeTrack algorithm still produces a reliable segmentation of sclera, lens and cornea. In order to determine whether the segmentation was of sufficient quality, we calculated the average energy functionals of sclera, lens and cornea for all participants (Table 1). Axial recordings were deemed unreliable if the functionals of cornea or lens were below 1.5, while this threshold was set at 0.5 for sagittal recordings. This excluded P1 and P3 from the analysis of axial recordings and P7 and P11 from sagittal recordings.

**Table 1:** Data quality assessment for each participant. Individual energy functional of sclera, cornea and lens as determined by normal gradient matching averaged over either all sagittally or axially acquired bSSFP scans (look into Kirchner et al. [13] for details about normal gradient matching).

|  |  | P1 | P2 | P3 | P4 | P5 | P6 | P7 | P8 | P9 | P10 | P11 |
| --- | --- | --- | --- | --- | --- | --- | --- | --- | --- | --- | --- | --- |
| E<sub>sclera</sub> | Axial | 2.47 | 2.17 | 2.26 | 2.26 | 2.25 | 2.31 | 2.30 | 2.42 | 2.43 | 2.35 | 2.42 |
|  | Sagittal | 2.13 | 2.15 | 2.10 | 2.04 | 2.63 | 2.17 | 1.68 | 2.09 | 2.08 | 2.02 | 1.73 |
| E<sub>cornea</sub> | Axial | 1.16 | 2.80 | 1.67 | 1.50 | 2.43 | 2.48 | 2.94 | 1.50 | 1.53 | 1.98 | 2.29 |
|  | Sagittal | 1.04 | 1.10 | 1.98 | 0.67 | 2.22 | 0.90 | 0.42 | 1.34 | 0.76 | 1.02 | 0.10 |
| E<sub>lens</sub> | Axial | 2.53 | 2.74 | 1.40 | 2.95 | 2.46 | 2.91 | 2.77 | 2.67 | 2.31 | 2.82 | 2.95 |
|  | Sagittal | 2.43 | 2.75 | 2.70 | 2.78 | 2.87 | 2.90 | 2.07 | 2.66 | 2.55 | 2.95 | 2.30 |

## Results

### Comparison of binocular retraction

To get an estimate of how precisely eyeball translation can be estimated from our data by the MREyeTrack algorithm, we compared anterior-posterior eyeball motion between both eyes from axial scans of the *Slow Blink* task. Throughout this study we refer to this motion as retraction, which is the posterior motion of the eyeball center during the fixation of the target dot. Since both eyes likely receive the same neural innervation during blinks, we expected that the two eyes exhibit the same translational motion. Indeed, we found high agreement between the measures of retraction between the two eyes for all participants (Fig. 1b,c & Movie 1 - https://doi.org/10.6084/m9.figshare.19732735.v1). Individual data points lie almost entirely on the identity line, indicating that both timing and amplitude of the retraction is almost identical between both eyes and that each eye receives the same neural innervation during blinks. The residuals from linear regression analysis had an average standard deviation of 0.12 mm, which is in agreement with modelling results from Kirchner et al. [13]. They claimed that eyeball translation could be estimated with a precision of 0.15 mm, based on comparing ground truth translation from artificial MR data with the estimated translation using the MREyeTrack algorithm.

**Figure 1:**
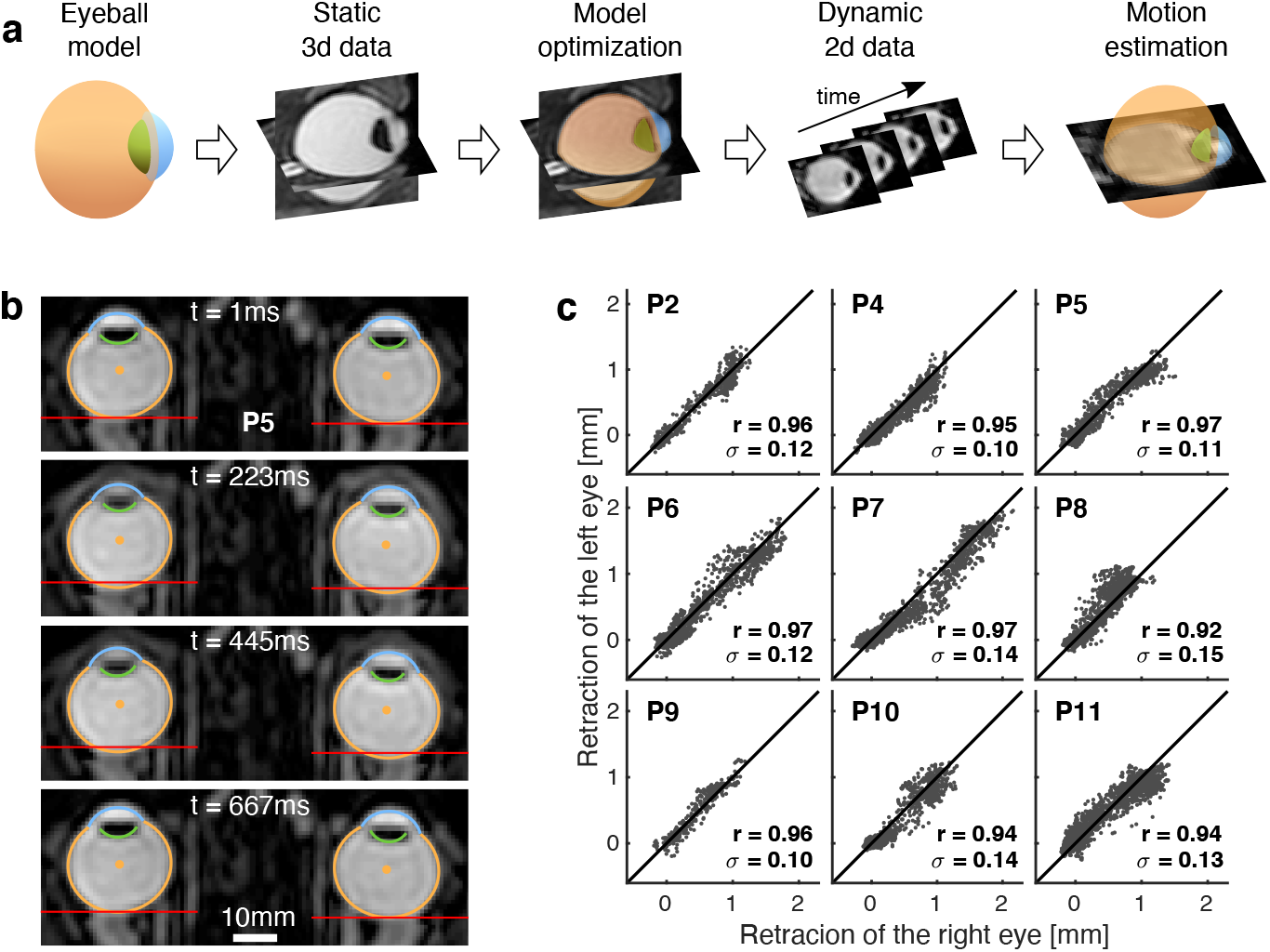
Estimation of eyeball motion using MREyeTrack. **a**, Illustration of the MREyeTrack workflow. An eyeball model consisting of sclera (orange), cornea (blue) and lens (green) is fitted to T2 weighted 3D data for each eye of each participant. Eyeball motion is then estimated by determining the optimal 2D projection of the eyeball model for each image of the dynamic MR scans. Reprinted from Kirchner et al. [13]. **b**, Axial recording of eyeball retraction during a blink of participant P5. Time is aligned to blink onset. The red lines are placed at the posterior eyeball border of the first image. The two middle images show simultaneous retraction of both eyes of around 1 mm. **c**, Comparison of retraction between left and right eye during the *Slow Blink* task for each participant. Each panel shows Pearson’s r and the standard deviation sigma of the residuals from linear regression analysis.

### Translational eyeball motion during blinks

We analysed eyeball translation along all three axes, i.e. translation from anterior to posterior which we called retraction, translation from inferior to superior which we called lifting and translation from medial to lateral. We observed only very little translational motion along the medial to temporal axis, which was typically in the range of 0.1 to 0.2 mm. No participant in any of the tasks showed medial or lateral translational motion of more than 0.5 mm. We focused the remainder of our analysis of translation on the measures of retraction and lifting in the sagittal plane. Retraction and lifting followed an identical time course, but lifting was often much larger than retraction (Fig. 2a & Movie 2 - https://doi.org/10.6084/m9.figshare.19732747.v1). For each participant, the ratio of lifting and retraction was constant for all blinks, meaning that blinks with a larger retraction also showed a proportionally larger lifting. Across participants, the ratio between lifting and retraction differed. For half of the participants the amount of lifting and retraction was roughly equal, while lifting was larger than retraction for the other half (Fig. 2b). Averaged across participants, the eyeball retracted by 0.79 mm (SD = 0.16 mm) and lifted by 1.35 mm (SD = 0.67 mm) during the *Slow Blink* task, showing that lifting is both larger and more variable than retraction. The maximum amount of overall translation that we measured throughout the experiment was 3.33 mm during a slow blink of P3.

**Figure 2:**
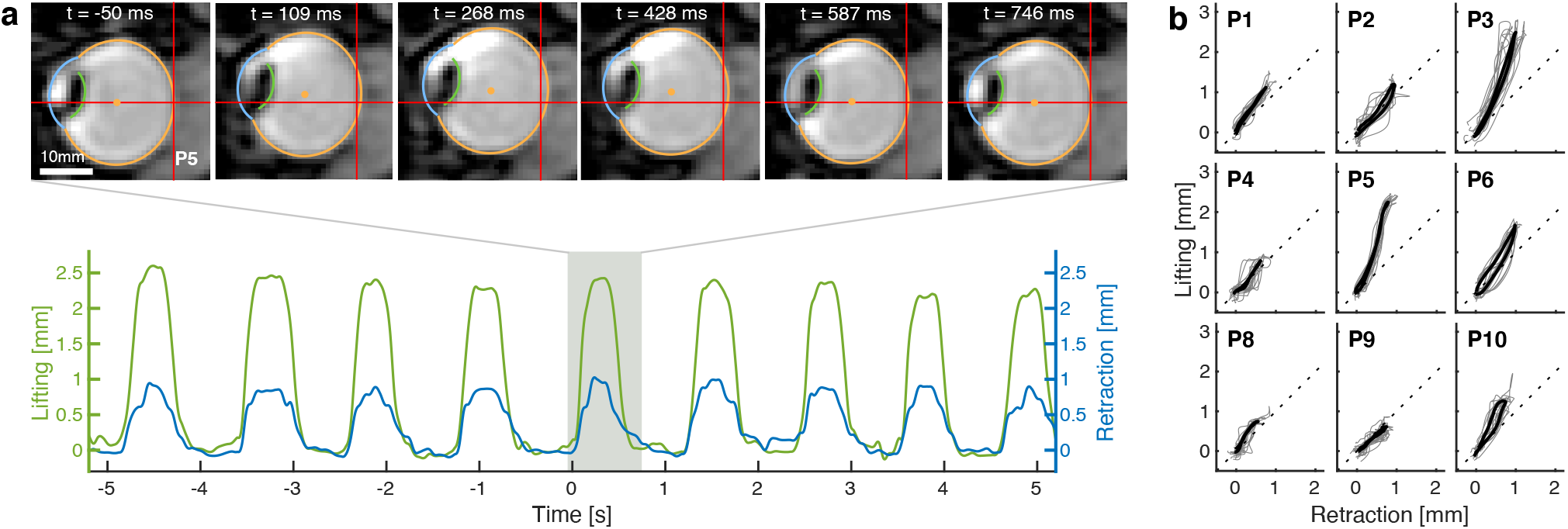
Eyeball lifting and retraction during blinks. **a**, Sagittal recording of eyeball lifting and retraction during the *Slow Blink* task of participant P5. Selected snapshots illustrate the translational eyeball motion with a red horizontal line placed at center position and a red vertical line at posterior border position of the first image. **b**, Translational eyeball trajectory during of each participant during the *Slow Blink* task. 10 randomly selected single trials are shown in grey, the average over all blink trajectories as a thick black line. Dotted line is the identity function. Note that the lifting component is at least as large as the retraction component and may even be twice as large.

### Holding state for prolonged lid closure

While the translational trajectories associated with blinks were fairly stereotypical when performing the same task, we often observed that amplitudes for the *Short Blink* task were smaller compared to the *Slow Blink* task. For even longer periods of lid closure during the *Eye Closure* task, there was no further increase in amplitude but instead the eyes held out in a retracted and lifted state while the eyes remained closed (Fig. 3a,b & Movie 3 - https://doi.org/10.6084/m9.figshare.19732750.v1). This holding state was typically reached after 200-300 ms, sometimes following a slight decay from full amplitude. Blinks performed during the *Short Blink* task often did not last long enough to reach this holding state, but nevertheless followed the same translational trajectory (Fig. 3c). In particular participants P3 & P6 showed a much smaller amplitude during short blinks, while their translational trajectory still overlapped with those of slow blinks and eye closures.

**Figure 3:**
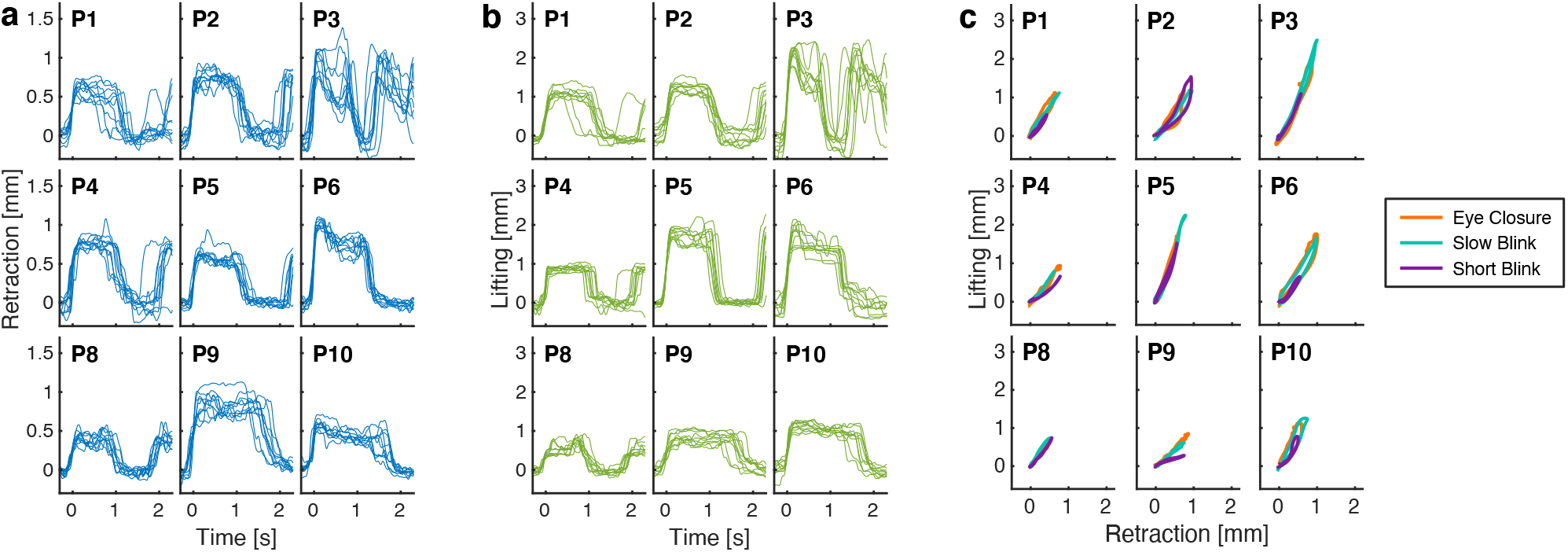
Holding state for prolonged eyelid closure. **a**, Retraction trajectory of 10 randomly selected trials from the *Eye Closure* task. After reaching full amplitude after around 200 milliseconds, the eyeball holds out in retracted location until the eyes are being opened again. **b**, The lifting component follows the same course as the retraction component and holds out after reaching full amplitude. Note the different scale compared to the retraction component. **c**, Averaged translational trajectories for each task of each participant. For many participants, the translation during short blinks does not reach full amplitude but follows the same path.

### Rotational eyeball motion during blinks

Rotations during short blinks in both axial and sagittal plane could be in either direction and could typically be described as monophasic motion with an amplitude in the range of 1-4°. This is in agreement with previous search coil studies, which investigated the rotational component of the blink-related eye movement in great detail [14, 15]. An interesting observation we made in our data from sagittal scans, was that some participants exhibited very large rotations for longer blinks. Most noticeably, P1 rotated upwards by up to 17° during the *Slow Blink* task (Fig. 4a & Movie 4 - https://doi.org/10.6084/m9.figshare.19732753.v1), but their rotations during short blinks were confined to 2° at maximum. These excessive rotations during slower blinks also lagged the onset of eyeball retraction by 100 ms and became larger with longer blink durations. Only some participants made these excessively large rotations which were also highly variable within participants and could be either up- or downwards (Fig. 4b).

**Figure 4:**
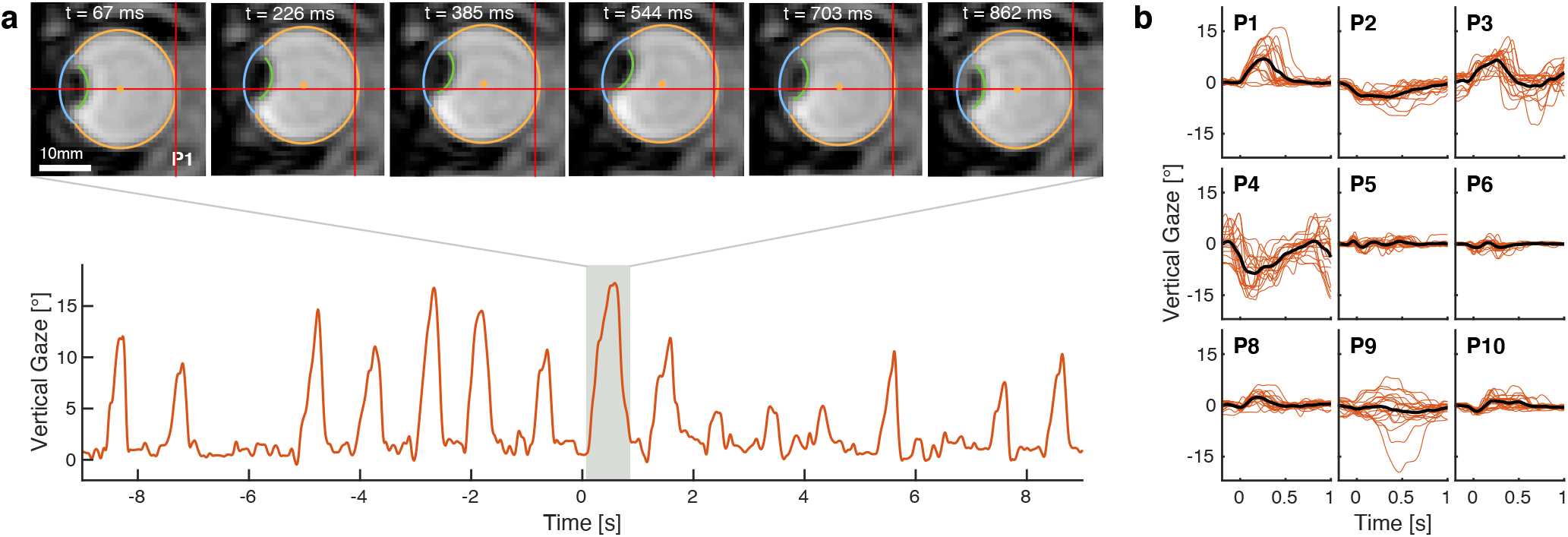
Eyeball rotation during blinks of long duration. **a**, Sagittal recording of vertical eyeball rotation during the *Slow Blink* task of participant P1. Selected snapshots show large rotational motion which scaled with blink duration. **b**, Rotational trajectories of 10 randomly selected blinks from the *Slow Blink* task for each participant as thin red lines and the averaged trajectory over all trials as thick black line.

### Eyeball compression

Since both origin and pulleys of the rectus muscles are located posterior to eyeball center, it is not unreasonable to assume that co-contraction of all 4 rectus muscles could compress the eyeball along the line of sight. Therefore, we calculated two measures of eyeball shape. The longitudinal diameter was defined as the distance from the border of the lens and vitreous body to the retina along the line of sight and through the eyeball center. The transversal diameter was defined to run perpendicular to the longitudinal diameter and passing through eyeball center (Fig. 5a). Only sagittal recordings were used for this analysis, because they contain less out-of-plane motion than axial recordings. Then, we tracked changes in diameter during the *Short Blink* task. We observed a sharp, consistent decrease in longitudinal diameter closely time-locked to blink onset for every single participant in the range of 0.30 to 0.85 mm (Fig. 5b). On average, the longitudinal diameter decreased by 0.59 mm (SD = 0.21 mm). In contrast to that, the transversal diameter remained constant for most participants and decreased only slightly for others (Fig. 5c). Even though the decrease in longitudinal diameter is small, it is visible in the MR footage by the posterior displacement of the border between lens and vitreous body relative to the segmentation (Fig. 5d & Movie 5 - https://doi.org/10.6084/m9.figshare.19732756.v1). Based on the video recordings, it was not clear to us whether this posterior lens movement could also be explained by an increase in lens diameter like during accommodation. In the video, it rather looks like the whole lens is moving posterior relative to eyeball center which would be consistent with eyeball compression. The anterior segment around the cornea is not well visible though, so the option of lens accommodation cannot be ruled out just by observing the video footage.

**Figure 5:**
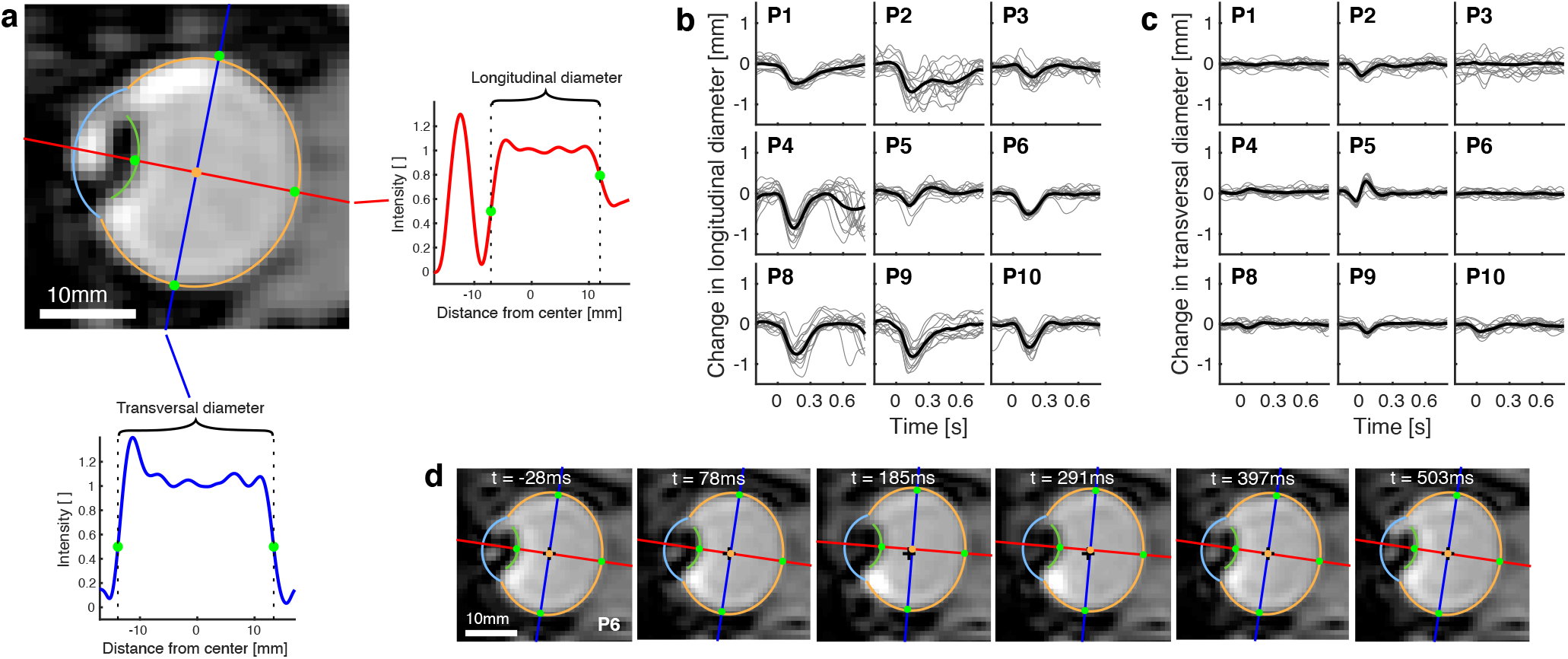
Eyeball compression during blinks. **a**, Illustration of calculation of longitudinal and transversal diameter based on image intensities relative to eyeball center. The orange marker is located at eyeball center, the green markers at eyeball borders. **b**, Change in longitudinal diameters during the *Short Blink* task, aligned to blink onset. 15 randomly selected trials as thin grey lines, average over all trials as thick black line. **c**, Same for change in transversal diameter. **d**, MR images of a short blink of participant P6. The black crosshairs mark eyeball center location in the first image. The green marker at the border between eyeball and lens moves posterior relative to lens segmentation in the two middle images.

## Discussion

Our results provide a detailed account of the full motion trajectory of blink-related eye movements. We demonstrated that the eyeball not only retracts but also lifts during a blink, and that the lifting component is on average almost twice as large as the retraction component. Only for blinks of longer duration does the translational motion reach its full amplitude and it then remains in a holding state for as long as the eyes are closed. For some participants, we also observed unusually large vertical rotations for blinks of longer durations, which lagged the onset of translational motion by a few dozen milliseconds. Furthermore, we investigated evidence for eyeball compression and found that the longitudinal diameter decreased by over half a millimeter while the transversal diameter remained roughly constant. This implies that the change in longitudinal diameter is not caused by out-of-plane motion of the eyeball, but is instead (we would argue) due to eyeball compression. These results show that the motion repertoire of eye movements is more complex than previously thought and raise some interesting questions regarding the innervation of extraocular muscles and orbital mechanics.

Blink-related eye movements were previously thought of as a combination of retraction and rotation. Our results show that eyeball lifting is an additional major component of the blink-related eye movement, arguably even its main component, and was found to occur consistently both during very short blinks and long periods of voluntary eye closure. This finding strengthens the hypothesis that the blink-related eye movement is primarily a translational motion due to co-contraction of extraocular muscles and that the accompanying rotational motion is an incidental derivative of the global translation [11, 14, 15]. In agreement with previous studies, we observed large tonic rotations during longer blinks but these were not only upwards but could be downwards as well [13, 14]. Large rotations during long or forceful blinks certainly contribute to the controversial reports regarding Bell’s phenomenon, but it is still puzzling why upwards rotation is predominantly observed during visual observation [16]. We think that the finding of eyeball lifting could explain the discrepancy between visual observation and modern eye tracking techniques. Lifting, like upwards rotation, leads to an elevation of the pupil which could lead to confusion of translational and rotational motion during visual observation where eyeball movement is deduced from movement of the pupil. For example, a 5° upwards rotation of an eyeball with a typical diameter of 26 mm elevates the pupil by 1.1 mm. This implies that a 2° downwards rotation paired with 1.4 mm of lifting would lead to an elevation of the pupil.

The observation of eyeball lifting also sheds new light on the neural innervation of extraocular muscles during blinks. It had been suggested that the co-contraction of inferior and superior rectus muscles would be sufficient to explain the blink-related eye movement [15], but the novel finding of eyeball lifting shows that other muscles must be involved as well. Our results suggest an activation of the superior oblique in order to explain eyeball lifting. It might be possible that there is an imbalance between superior oblique and its antagonist the inferior oblique, but this might not even be necessary. The superior oblique is unique among the six eye muscle in that its tendon is guided through the trochlea before innervating the eyeball. The other eye muscles have soft tissue pulleys which actively move when the muscle contracts [29, 30]. The pulley of the superior oblique however is the rigid trochlea, which produces additional leverage on the eyeball from the superior oblique presumably leading to an eyeball lift. Additional evidence for an active role of the superior oblique in eyeball lifting comes from the analysis of global eyeball position as a function of gaze direction [18]. It was found that downwards gaze, which is accomplished by the contraction of inferior rectus and superior oblique, lifts up the eyeball. In contrast, the contraction of superior rectus and inferior oblique produces upwards gaze but lowers the eyeball as a whole. Interestingly, EMG recordings in rabbits have shown that all extraocular muscle except the superior oblique contract during blinks [11]. Rabbits, like many mammals, have a retractor bulbi, an eye muscle which is specifically responsible for eyeball retraction. Perhaps the superior oblique has taken over this task in humans.

Our results further suggest that simultaneous contraction of the rectus muscles lead to a slight compression of the eyeball. Based on the video recordings, we considered whether the decrease in longitudinal diameter really reflects eyeball deformation or instead could be explained by lens accommodation. However, we are not aware of any reports on accommodation during blinks and an earlier MRI study showed that the lens diameter increased by only 0.33 mm on average during accommodation [31]. This is too small to explain the 0.59 mm decrease in longitudinal diameter that we observed. Additionally, it seems plausible to us that the enlarged force applied by the eye muscles during blinks is powerful enough to result in a slight deformation. In this context, it would be interesting to investigate whether the amount of force exerted on the eyeball by compression and translation could have long term effects on the eyeball shape and might even be related to certain pathologies. There are other examples in the literature where mechanical force being exerted on the eye has been suggested have long term consequences. For example, the force exerted by eye rubbing has been hypothesised to be the cause of keratoconus [32].

Dynamic MRI proved to be a valuable tool to study ocular motility during blinks. We obtained detailed trajectories of this motion using the MREyeTrack algorithm. We found that the blink-related eye movement could be accompanied by translations of up to 3.3 mm, far exceeding the extent previously reported. Given the frequency of blinks in normal life, these translations should be considered a fundamental part of oculomotor control. To the best of our knowledge, translations of this magnitude have so far not been incorporated in biomechanical modelling of the oculomotor plant. Therefore, blink-related eye movements could be used to test the validity of different models. Future studies could focus on the effect of blinking and the affiliated co-contraction on extraocular muscles and the pulley system.

## Data availability

All data are available from the corresponding author upon reasonable request.

## Acknowledgments

This work has received funding from the European Union’s Horizon 2020 research and innovation programme under the Marie Skodowska-Curie grant agreement No. 734227.

